# Vectorial boundary representation of 3D layout in human visual cortex

**DOI:** 10.1101/2025.11.17.688791

**Authors:** Yichen Wu, Sheng Li

## Abstract

Human spatial navigation relies on the brain’s ability to visually represent the 3D layout of the environment. To understand how the brain encodes the layout information, it is crucial to identify the key features of environmental layout and how they are processed in the human brain. The vector coding principle, which highlight the role of boundary distance and orientation, provides a theoretical framework supported by physiological evidence from rodents. In this study, we developed a reconstruction approach to quantitatively estimate 3D layout information from natural indoor scene images. This approach enabled analyses of fMRI data from the large-scale Natural Scenes Dataset (NSD) using vector-based models of 3D layout. To validate the NSD-based results and examine task-related dynamics, we further conducted fMRI and MEG experiments with navigation-related and non-navigational tasks. Controlling for low-, mid-, and high-level visual and semantic features of natural indoor scenes, we found a spatiotemporal dissociation between boundary distance and orientation representations in the human brain. Relative distance was encoded in the early visual cortex during early processing in a task-invariant manner, whereas orientation was represented in scene-selective higher visual areas during later processing and was modulated by navigation-related tasks. Importantly, task modulation manifested as feedback-induced enhancement of orientation coding in the early visual cortex. Together, these findings provide a novel perspective on how the human brain represents navigation-relevant information about the immediate surrounding environment, advancing our understanding of the neural mechanisms that link perception to action in spatial navigation.

## Introduction

Spatial navigation depends on the brain’s ability to explore and represent the external world. Seminal discoveries in rodents, particularly the identification of place cells (O’Keefe & Dostrovsky, 1971) and grid cells (Hafting et al., 2005), have laid the foundation for understanding the neural mechanisms underlying spatial map construction. In humans, navigation relies predominantly on vision to represent the complex three-dimensional (3D) world, especially the spatial geometry of surrounding environment (Spelke & Lee, 2012). This visual representation of environmental layout is essential for reconstructing egocentric boundaries, which in turn form the basis for constructing allocentric cognitive maps (Peer & Epstein, 2021). In the human brain, a network of scene-selective areas, including the occipital place area (OPA), parahippocampal place area (PPA), and retrosplenial complex (RSC), has been identified as crucial for representing the local environment. These regions respond more strongly to images of landmarks, buildings, and rooms than to faces or objects, underscoring their central role in processing structural layout information (Epstein & Baker, 2019).

Boundaries define the limits of exploration within an environment, making them critical elements of spatial representation in the brain. In a given space, walls serve as invariant components that determine the overall layout, whereas most objects are movable and subject to change. Thus, the arrangement of walls in a scene can be regarded as a primary representation of spatial layout. Previous studies on human spatial cognition have shown that scene-selective areas, particularly the OPA, are activated by walls (Epstein & Kanwisher, 1998; Ferrara & Park, 2016; Kamps et al., 2016) and encode navigationally relevant information (Bonner & Epstein, 2017; Harel et al., 2013; Kravitz et al., 2011; J. Park & Park, 2020; S. Park et al., 2011; Persichetti & Dilks, 2018). More recent work using artificial stimuli to model 3D geometries further demonstrated that these areas also encode scene layout (Henriksson et al., 2019; Lescroart & Gallant, 2019). Notably, Lescroart & Gallant (2019) found that scene-selective areas are tuned to the 3D configuration of surfaces in artificial scenes, even after controlling for low-level 2D visual features. This surface-based coding model provide valuable insights into how the human brain visually represents navigational information. However, surface-based models do not capture boundary vectors, a critical component of spatial representation identified in rodent navigation systems (Bicanski & Burgess, 2020).

Successful navigation requires local representation of boundary information in an egocentric reference frame to construct the layout of agent’s surroundings (e.g., front, back, left, and right). According to the principle of vector coding, neural representations of environmental features are organized by the direction and distance of boundaries relative to the agent, which together constitute the fundamental structure of spatial layout (Bicanski & Burgess, 2020). In rodents, neurons with vector-based receptive fields have been identified for boundary (Alexander et al., 2020; Hinman et al., 2019) and border (Barry et al., 2006; Lever et al., 2009; Solstad et al., 2008), underscoring the essential roles of boundary vectors representations in spatial navigation. Despite these findings, how such vector-based coding is implemented in the human brain remains largely unknown.

Given the complexity with which the human brain transforms 2D natural images into 3D layout representations, large-scale neuroimaging datasets are particularly valuable for investigating these representations under naturalistic conditions. A growing trend in functional Magnetic Resonance Imaging (fMRI) research involves intensive scanning, which enables detailed analyses of large datasets from individual participants (Kupers et al., 2024). A prominent example is the Natural Scenes Dataset (NSD), which adopts this approach with a recognition task using natural images (Allen et al., 2022). The NSD provides a unique opportunity to study how the human brain represents spatial layout. However, unlike artificial stimuli, the natural images in the NSD lack ground-truth 3D layout annotations, limiting their use in fine-grained analyses of spatial cognition.

To address this limitation, we developed a computer vision-based approach to reconstruct 3D scene layouts from natural indoor images. Applying this method to the NSD enabled us to examine fine-grained neural representations of spatial layout in this large-scale 7T fMRI dataset. Because the NSD involved a non-navigational task, we further conducted two complementary experiments, using fMRI and magnetoencephalography (MEG), to validate and extend the NSD-based findings. These experiments incorporated both a navigation-related layout discrimination task and a non-navigational texture discrimination task, using naturalistic indoor images with ground-truth maps of boundary distance and orientation derived from the Matterport3D database. Across these datasets, we found converging evidence for a spatiotemporal dissociation between distance and orientation representations of layout boundaries, as well as task-dependent modulation in navigation-related context.

## Results

### A new approach for reconstructing 3D layout and self-pose from natural indoor images

Investigating how the brain represents 3D layout in large-scale natural image datasets (e.g., NSD) has been challenging because explicit layout annotations are typically available only for artificial stimuli. To overcome this limitation, we developed a reconstruction approach that quantitatively estimates both the 3D layout and self-pose of natural indoor images through an annotation-based procedure (Figure 1A, see Methods for details). Our approach uses a vanishing point-based camera calibration technique, leveraging the fact that parallel lines in the 3D world converge within the 2D projection of a scene. Six human annotators manually identified six key edges in each image. Assuming that the optical point coincides with the image center, we recovered essential camera parameters, including focal length, field of view, pitch, roll, and wall orientations. Based on these parameters, we generated 3D layout segmentation maps for 2,120 NSD indoor images (see Figure 1B for reconstruction examples).

**Figure 1:**
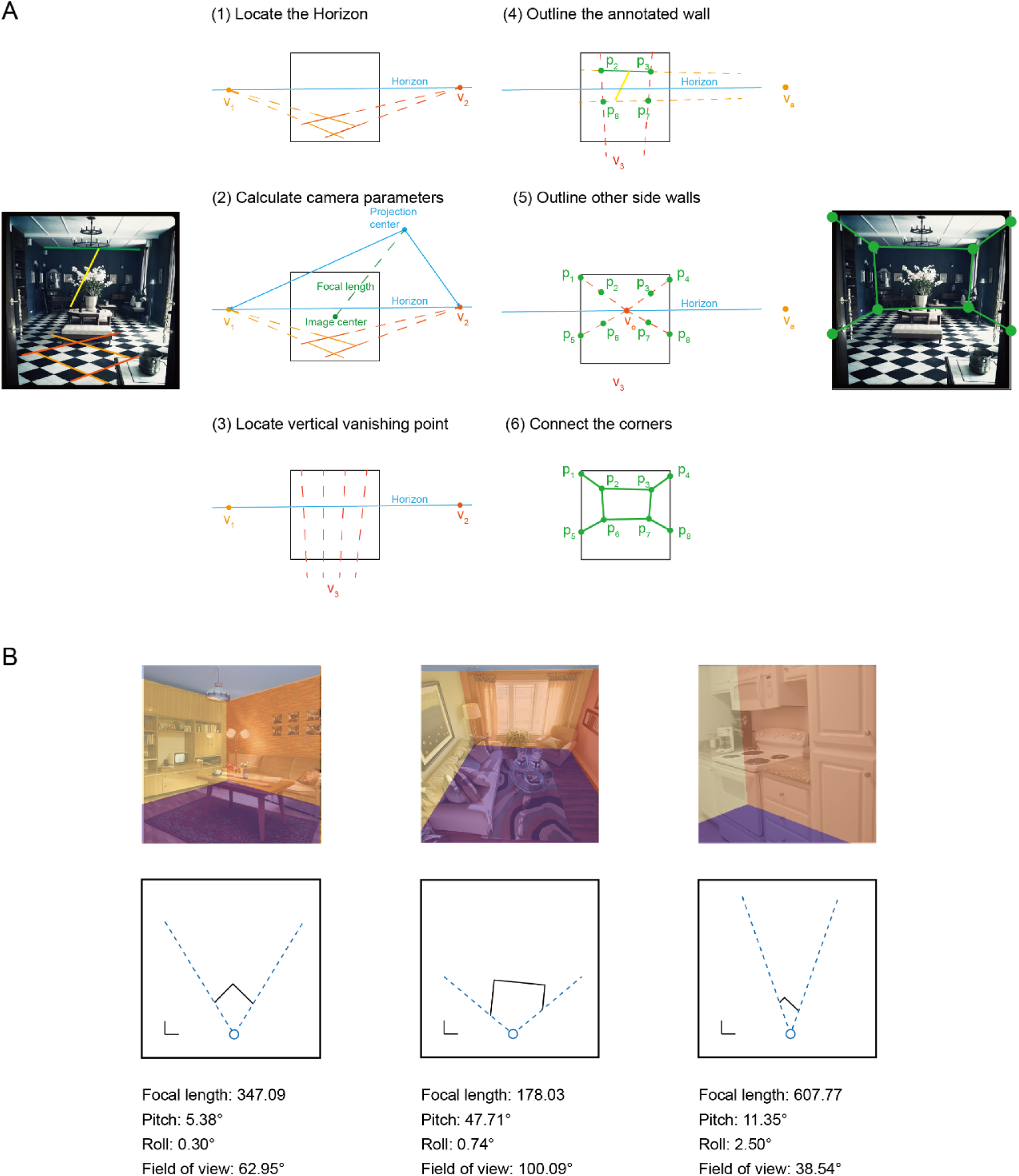
Reconstruction of 3D layout and self-pose for natural indoor images. **A**, Schematic of the reconstruction approach for an example image (see Methods for details). The left image shows the six manually labelled edges: the two pairs of orange line are the horizontal edges parallel to the ground, while the green line and yellow line were used to calculate the height and width of the middle wall, respectively. The middle panel illustrates the reconstruction procedure based on the labelled lines. The right image shows the reconstructed corners and wall conjunctions. **B**, Examples of reconstruction results. The top row displays three NSD indoor images overlaid with reconstructed layout segmentation maps. The middle row presents the aerial-view maps with side walls marked by black lines. The blue circle indicates the observer, and the dashed blue lines define the field of view. Reconstructed parameters are shown at the bottom.

To evaluated the robustness of our method, we tested on the Matterport3D database, which provides ground-truth camera parameters and layout annotations. Across 40 undistorted indoor images, the mean errors in pitch and roll angles were 1.65° (SD = 2.38°) and 1.13° (SD = 1.28°), respectively. Layout reconstruction accuracy, measured by the proportion of misclassified pixels relative to the ground-truth segmentation map, yielded a mean pixel error was 2.98% (SD = 3.64%). Because NSD stimuli were not distortion-corrected, the reconstruction accuracy is slightly reduced compared with the Matterport3D images, as the annotated lines that are supposed to be parallel to the ground may contain errors.

### Dissociated representations of boundary distance and orientation in the visual cortex: NSD experiment

The positions of side walls define the boundaries of an indoor space and are essential for ground-level navigation. From an egocentric aerial view, side walls can be modeled as orthogonal lines radiating from the observer (Figure 1B), a geometric structure that lies at the core of vector coding principle (Bicanski & Burgess, 2020). While our reconstruction approach estimates the orientations of side walls based on their intersection with the horizon, it does not directly recover precise distance. We therefore approximated the relative distance of side walls by computing the area difference between side wall regions and ceiling/floor regions. These reconstructed parameters were then used to generate layout models for walls’ relative distance and orientation (Figure 2C).

**Figure 2:**
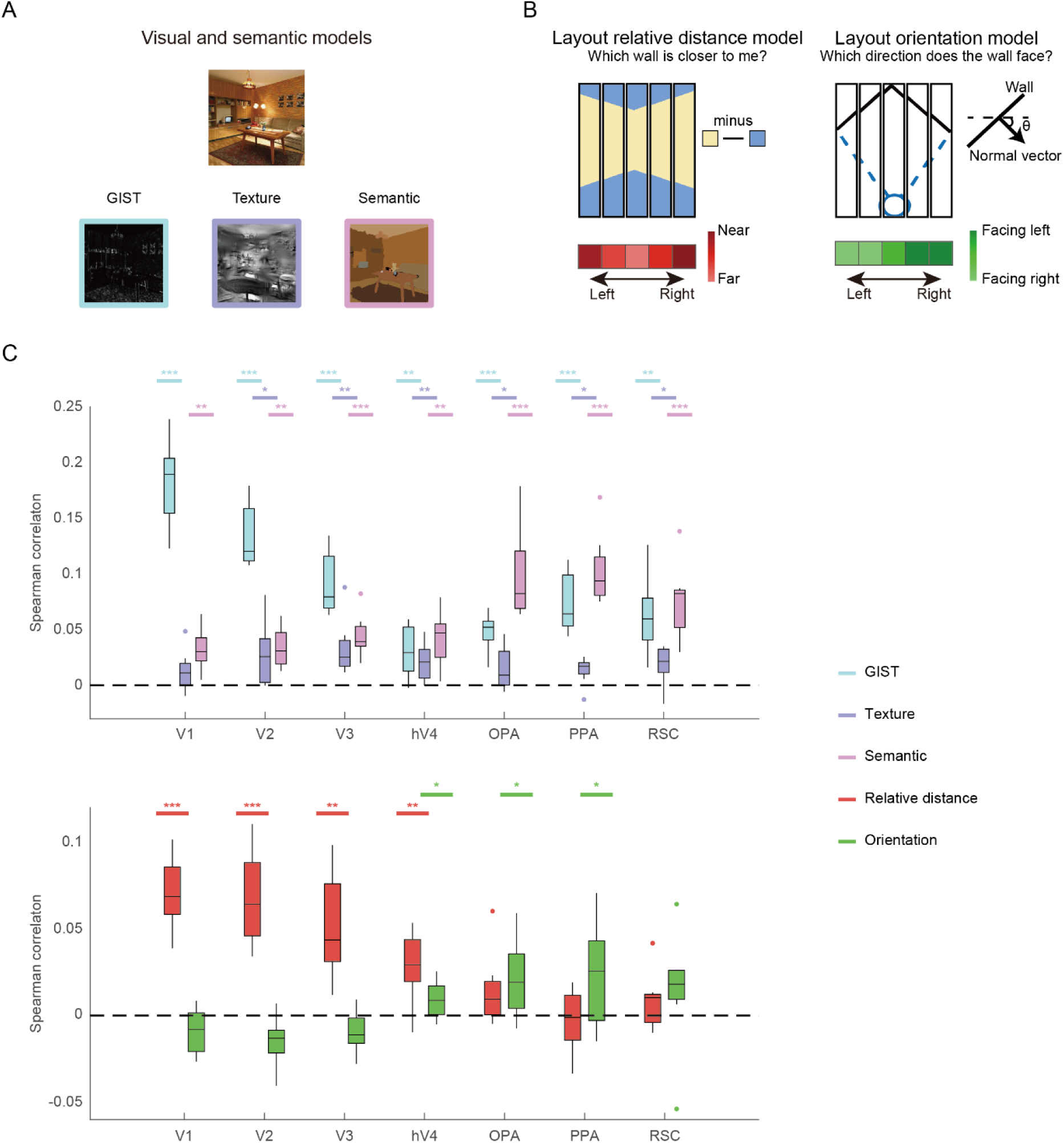
Model representations for 2D features and 3D layout: NSD experiment. **A**, Models for 2D visual and semantic features. **B**, Models for 3D layout features, including boundary relative distance and orientation. **C**, RSA results across ROIs. The upper plot shows the partial correlations between 2D model RDMs and neural RDMs. The lower plot shows the partial correlations between 3D model RDMs and neural RDMs. Both sets of results were derived from the same partial correlation analysis but are displayed separately for clarity. Asterisks indicate significance in one-tailed t-test against chance level (0). * *q* < 0.05; ** *q* < 0.01, *** *q* < 0.001.

Because layout features covary with 2D image statistics, we incorporated three additional models to control for low-, mid-, and high-level visual and semantic features (Figure 2A): the GIST model for low-level features of spatial frequency and orientation (Oliva & Torralba, 2001), the S-P model for mid-level features of texture (Portilla & Simoncelli, 2000), and the object2vec model for high-level features of semantic and object cooccurrence (Bonner & Epstein, 2021). We constructed representational dissimilarity matrices (RDMs) for each model and region of interest (ROI). We then conducted representational similarity analysis (RSA) using partial Spearman correlation to assess the unique contribution of each model while controlling for the others.

We found that 2D visual and semantic features are significantly partial correlated with neural activities across both early visual and scene-selective areas (Figure 2B). Specifically, correspondence between the GIST, texture, and semantic models and low-, mid-, and high-level visual regions confirmed that these models capture the intended representational hierarchy. Critically, after controlling for these 2D visual and semantic features, early visual areas exhibited significant partial correlations with the relative distance model (V1: t(7) = 9.72, *q* < 0.001; V2: t(7) = 7.05, *q* < 0.001; V3: t(7) = 4.95, *q* = 0.002; hV4: t(7) = 3.94, *q* = 0.006; one-tailed, FDR corrected), whereas hV4, OPA and PPA showed significant partial correlations with the orientation model (hV4: t(7) = 2.50, *q* = 0.029; OPA: t(7) = 2.67, *q* = 0.023; PPA: t(7) = 2.20, *q* = 0.042; Figure 2D).

These results revealed a clear dissociation along the visual hierarchy: the early visual cortex predominantly represents boundary distance, whereas scene-selective regions encode boundary orientation. To confirm this dissociation at the individual-subject level, we performed a searchlight analysis within the visual cortex of each participant. Figure 3 illustrates an example participant’s flattened cortical surface maps showing the distributions of correlation for the relative distance and orientation models after controlling for 2D visual and semantic features (see Supplementary Figure 3 for maps of other NSD participants). The surface map demonstrates a posterior-anterior gradient from distance to orientation representation: voxels in the posterior occipital cortex uniquely correlated with the relative distance model, while anterior occipital and temporal regions showed stronger correlations with the orientation model. Importantly, these effects could not be accounted for by alternative 3D layout models, such as mean depth model or the models used in Henriksson et al. (2019) and Lescroart & Gallant (2019); both boundary relative distance and orientation models remained significant after controlling for these alternatives (Supplementary Figure 2). Together, these analyses indicate distinct and hierarchically organized neural representations of boundary distance and orientation along the human visual processing stream.

**Figure 3:**
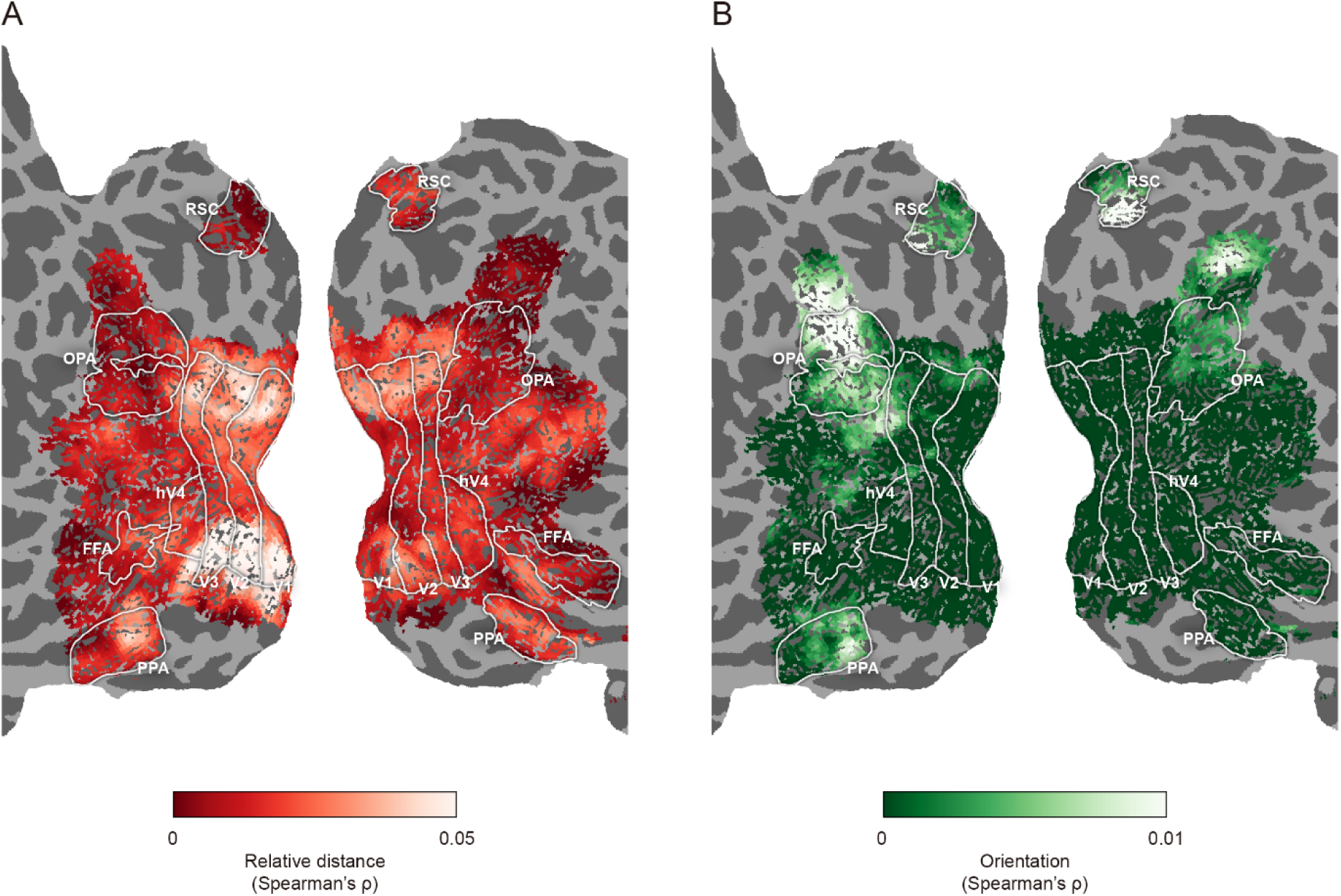
Cortical representations of 3D layout for one example participant: NSD experiment (Subject 1 in NSD dataset). Searchlight-based partial correlations are shown on flattened cortical surface maps (see Supplementary Figure 3 for maps of other participants). **A**, Representation of the relative distance model. **B**, Representation of the orientation model.

### Representation of self-pose in the visual cortex: NSD experiment

Self-pose parameters such as pitch and roll angles influences how the visual system interprets environmental structure, yet direct neural evidence for their encoding in the human visual cortex has remained limited. Using the reconstructed pitch and roll angles of NSD indoor images, we tested whether self-pose information could be decoded from the fMRI responses in early visual and scene-selective regions (Figure 4).

**Figure 4:**
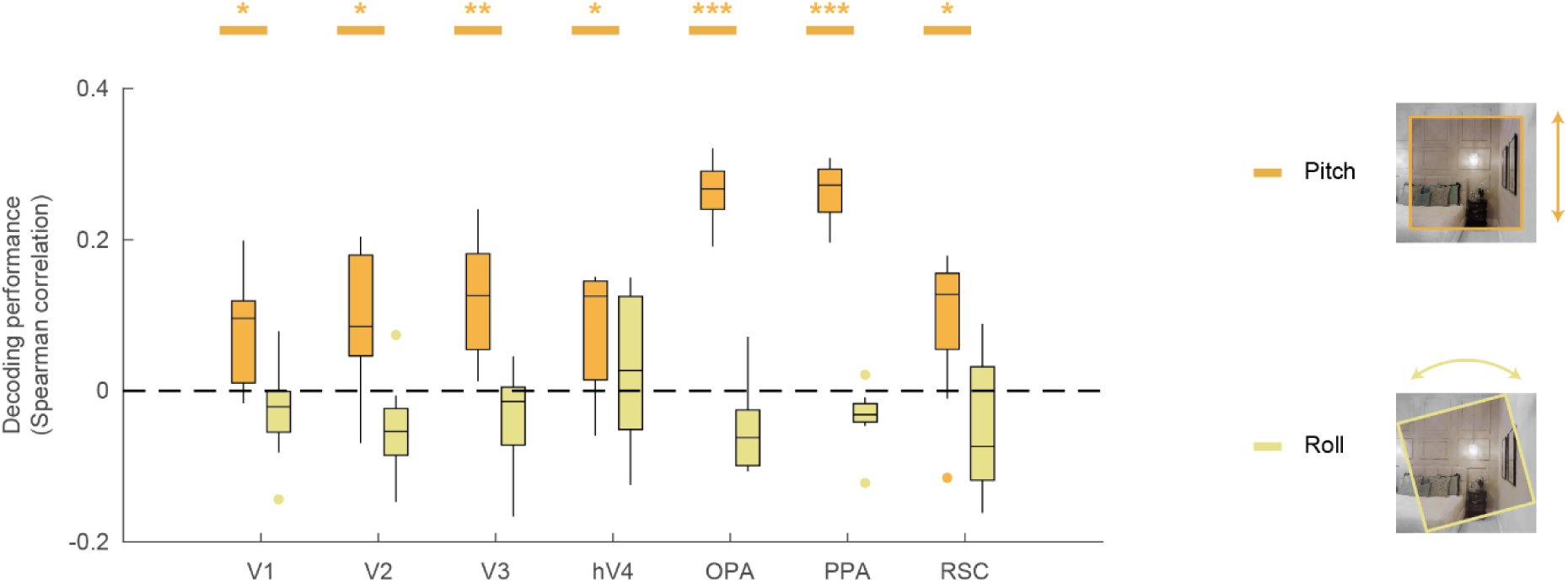
Decoding self-pose from fMRI signals: NSD experiment. Pitch and roll angles were decoded using support vector regression (SVR). Only pitch angle showed reliable decoding across the visual cortex. Asterisks indicate significance in one-tailed t-test against chance level (0). * *q* < 0.05; ** *q* < 0.01, *** *q* < 0.001.

Significant prediction of pitch angles was observed across all early visual and scene-selective regions (V1: t(7) = 3.08, *q* = 0.028; V2: t(7) = 2.91, *q* = 0.028; V3: t(7) = 4.42, *q* = 0.007; hV4: t(7) = 2.87, *q* = 0.028; OPA: t(7) = 18.52, *q* < 0.001; PPA: t(7) = 18.52, *q* < 0.001; RSC: t(7) = 2.57, *q* = 0.037). In contrast, roll angles could not be reliably decoded in any ROI, indicating that pitch, but not roll, is encoded in the visual cortex. This finding provides direct evidence that the visual system contains representations of navigation-relevant self-pose information, consistent with the behavioral importance of pitch during vertical navigation (e.g., ascending or descending stairs).

### Task-dependent enhancement of layout representation in early visual cortex: fMRI experiment

The NSD analyses revealed that both early visual and scene-selective areas encode layout and self-pose information even when scene layout is task-irrelevant. To determine how these representations are modulated when layout becomes behaviorally relevant, we conducted a new fMRI experiment. Participants performed two block-designed tasks: a layout discrimination task, which required active encoding of the spatial layout, and a texture discrimination task, which emphasized surface texture and color and served as a non-navigational control (Figure 5A).

**Figure 5:**
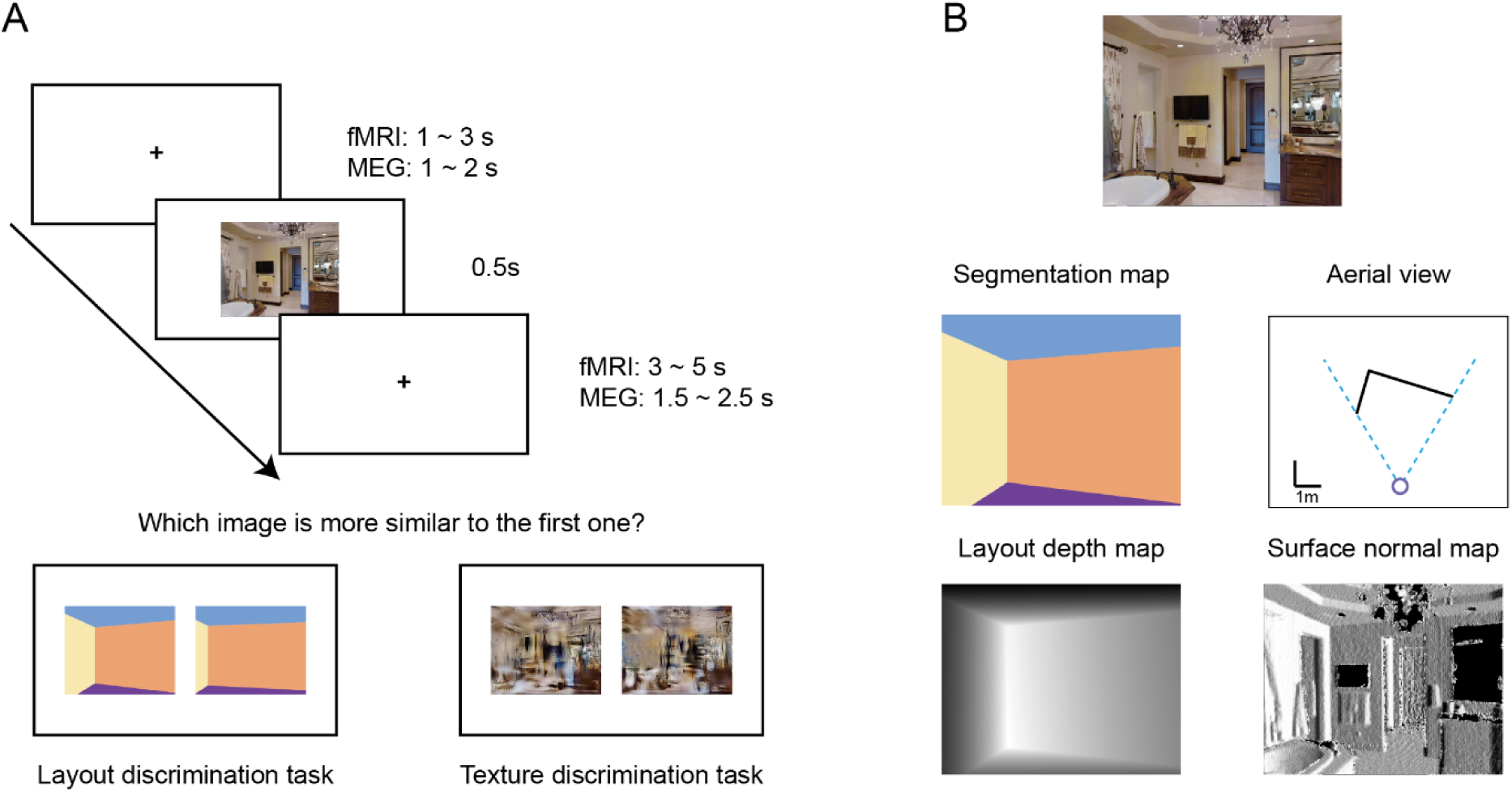
Task procedures for the Matterport3D fMRI and MEG experiments. **A**, Match-to-sample paradigms for layout discrimination and texture discrimination tasks. Inter-trial and inter-stimulus intervals are adjusted to meet of fMRI and MEG requirements. **B**, Example scene image from the Matterport3D database and its ground-truth layout-related annotations.

Stimuli were drawn from the Matterport3D database (Chang et al., 2017), which provides ground-truth camera poses as well as depth, segmentation, and surface normal maps (Figure 5B). These annotations enabled precise construction of distance and orientation models for side walls. To avoid potential confounds from self-pose variation, all images were selected with pitch and roll angles fixed at 0°. Task difficulty was equated through a pilot experiment, with accuracies of 77.60% and 76.32% for layout and texture tasks, respectively (t(29) = 0.41, *p* = 0.683).

We performed RSA on pre-defined ROIs, including V1 and scene-selective areas (OPA, PPA, and RSC). Figure 6 shows partial correlations between neural RDMs and the layout models after controlling for 2D visual and semantic features. We observed robust task-dependent modulation of layout representations. In the layout discrimination task (Figure 6A, right), both V1 and OPA showed significant partial correlations with relative distance (V1: t(29) = 8.41, *q* < 0.001; OPA: t(29) = 2.99, *q* = 0.011) and orientation (V1: t(29) = 5.61, *q* < 0.001; OPA: t(29) = 3.88, *q* = 0.001) models. In contrast, in the texture discrimination task (Figure 6B, right), V1 and PPA showed significant partial correlations with the relative distance model (V1: t(29) = 4.68, *q* < 0.001; PPA: t(29) = 2.71, *q* = 0.016), while only OPA showed a significant partial correlation with orientation (t(29) = 2.78, *q* = 0.015). Crucially, direct task comparison revealed significantly enhanced representation in V1 for both relative distance (t(29) = 3.99, *q* = 0.002, two-tailed, FDR corrected) and orientation (t(29) = 5.21, *q* < 0.001) models (Figure 6C).

**Figure 6:**
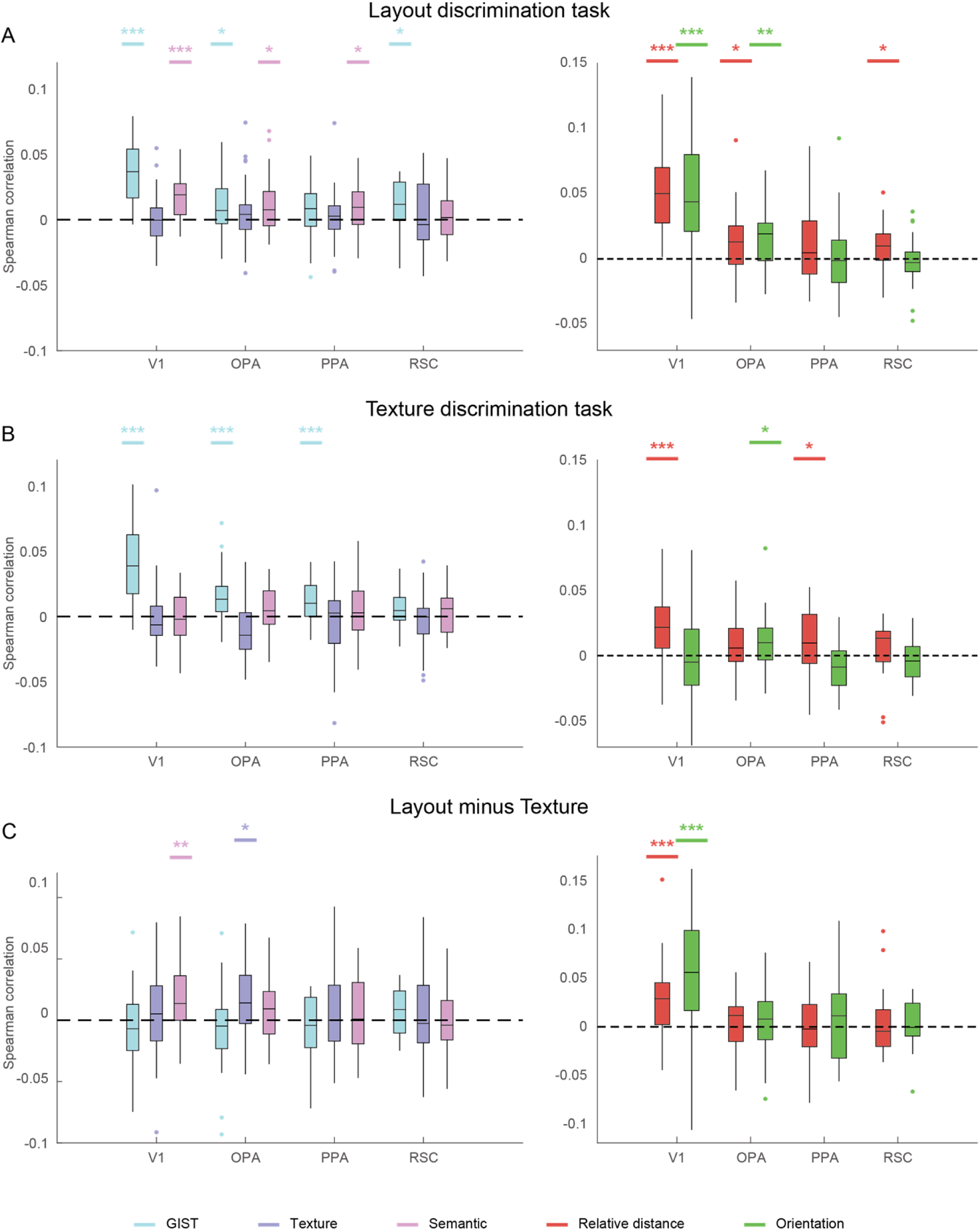
Model representations for 2D features and 3D layout: Matterport3D fMRI experiment. The left plots show partial correlations between 2D model RDMs and neural RDMs (see Supplementary Results for statistics). The right plots show partial correlations between 3D layout model RDMs and neural RDMs. **A**, Layout discrimination task. **B**, Texture discrimination task. **C**, Task-dependent enhancement of representation, calculated as the difference between partial correlation coefficients for the two tasks (layout minus texture). Asterisks in **A** and **B** denote significant results in one-tailed t-test against chance (0), while asterisks in **C** indicate significance based on two-tailed tests. * *q* < 0.05; ** *q* < 0.01, *** *q* < 0.001.

To further examine whether the brain encodes metric depth information, we built a precise distance model using the ground-truth depth maps from Matterport3D. Neither this precise distance model nor a mean depth model (Supplementary Methods) showed significant correlation with neural activity (Supplementary Figure 4), suggesting that precise metric distance computation may require additional reference objects and longer processing time.

Together, these results highlight two key implications. First, the texture discrimination task replicates the NSD-based findings, supporting task-invariant encoding of spatial layout in the visual cortex. Second, the layout discrimination task reveals enhanced representations of boundary distance and orientation when layout information becomes task-relevant, suggesting possible feedback modulation in V1. However, the limited temporal resolution of fMRI precludes direct characterization of such feedback processes, motivating a subsequent MEG experiment.

### Temporal dissociation of distance and orientation representations: MEG experiment

To characterize the temporal dynamics of layout representations and their task-dependent modulation, we conducted an MEG experiment using the same stimuli and design as in the fMRI experiment. Participants achieved comparable accuracies across tasks (layout: 72.73%; texture: 72.52%; t(31) = 0.18, *p* = 0.856).

For each task, we constructed a neural RDM at each time point of the MEG signal using occipital channels. RSAs were performed between neural RDMs and model RDMs. Partial correlation analysis revealed significant temporal clusters correlated with the GIST and semantic models in the layout discrimination task (*p* < 0.05, TFCE one tailed; Figure 7A, left) and with the GIST model along in the texture discrimination task (*p* < 0.05; Figure 7B, left).

**Figure 7:**
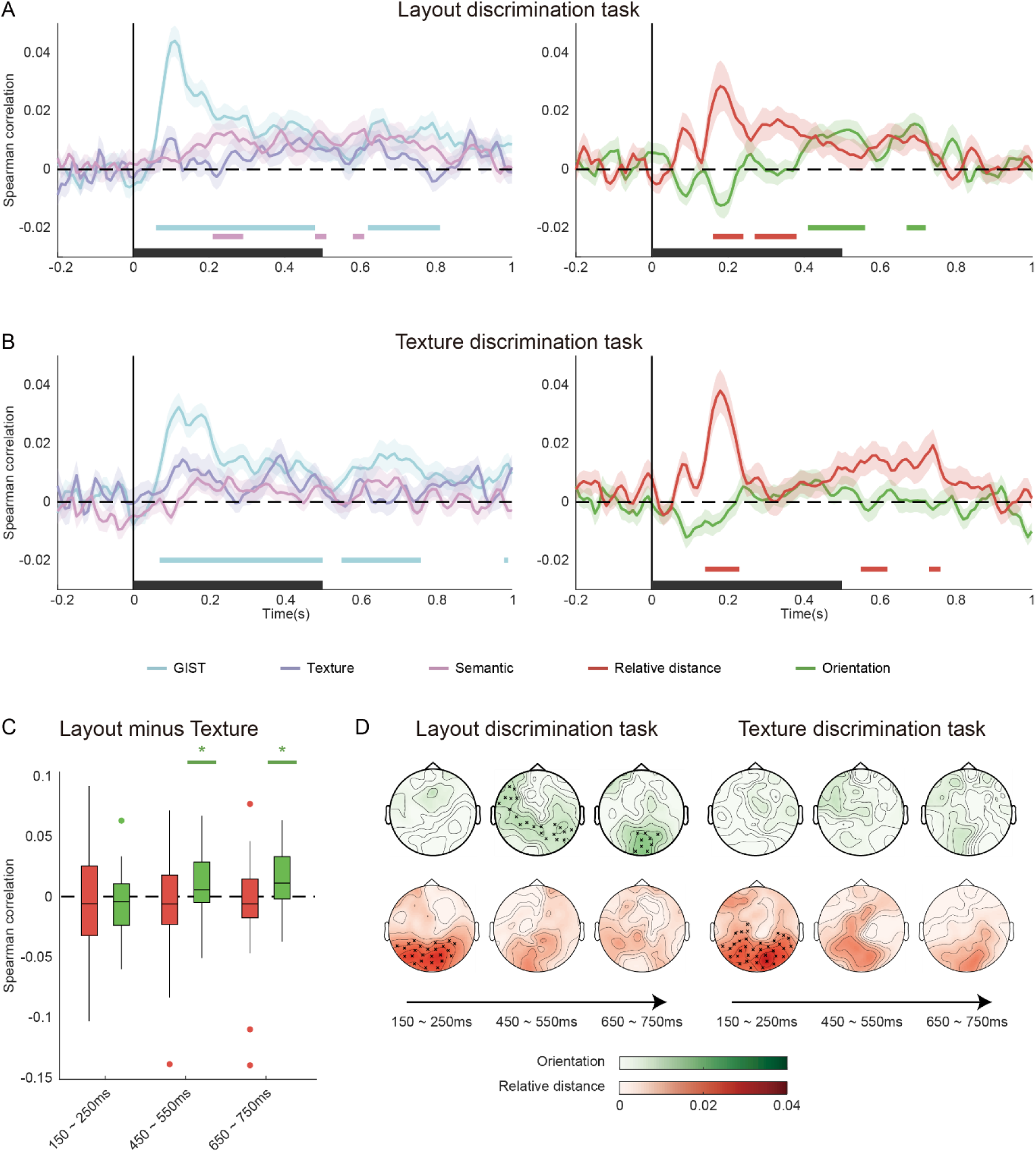
Model representations for 3D layout: Matterport3D MEG experiment. **A**, Layout discrimination task. **B**, Texture discrimination task. For **A** and **B**, significance was determined using one-sample t-tests against chance level, corrected for multiple comparisons with TFCE one-tailed *p* < 0.05. Colored horizontal bars indicate significant clusters, and black bars on the x-axis denote image presentation periods. **C**, Task-dependent enhancement of layout representation in three time windows. Asterisks denote significance in one-tailed t-test against chance level. * *q* < 0.05; ** *q* < 0.01, *** *q* < 0.001. **D**, Searchlight results. Scalp maps show partial correlations of the relative distance (red) and orientation (green) model RDMs with neural RDMs. Crosses mark significant clusters (TFCE one-tailed, *p* < 0.05).

Critically, relative distance representations emerged in both tasks (Figure 7A and 7B, right): 160-240 ms and 270-380 ms for the layout discrimination task, and 140-230 ms, 550-620 ms, and 730-760 ms for the texture discrimination task. In contrast, orientation representations appeared exclusively in the layout discrimination task within later time windows (410-560 ms and 670-720 ms), demonstrating a temporal dissociation between distance and orientation coding.

To quantify task-dependent effects, we focused on three 100-ms windows centered at 200 ms, 500 ms, and 700 ms (Figure 7C), which corresponded to time periods showing significant RSA correlations with the layout models. Task-dependent enhancement was evident at 500 ms and 700 ms windows (450-550 ms: t(31) = 2.85, *q* = 0.023; 650-750 ms: t(31) = 3.06, *q* = 0.023; two-tailed, FDR corrected). Searchlight analysis across channels corroborated these results (Figure 7D): early clusters (∼200 ms) over posterior channels correlated with relative distance in both tasks, while orientation-related clusters (∼500 and 700 ms) appeared only in the layout discrimination task, particularly around occipital channels at 700 ms.

Collectively, these findings reveal a clear temporal dissociation in MEG responses, with early distance coding followed by later orientation coding that parallels the spatial dissociation observed in fMRI. This temporal sequence suggests a hierarchical and dynamic organization of boundary feature representations in the human brain. At the same time, the delayed emergence of orientation coding aligns with known feedback processing windows (Brandman & Peelen, 2023; Groen et al., 2016; Kaiser et al., 2019), implying that feedback-related processes may further refine orientation representations at later stages.

### Linking fMRI and MEG responses

To directly relate the spatial and temporal findings, we compared representational patterns across modalities. Group-averaged RDMs from V1 and three scene-selective areas in the fMRI experiment were partial correlated with MEG RDMs from occipital channels at each time point, without incorporating any feature-based model RDMs. This analysis aimed to identify shared representational patterns between hemodynamic and electromagnetic signals independent of specific encoding models.

The results showed that fMRI responses in V1 made a significant and sustained contribution to MEG responses in occipital channels across both tasks. In the layout discrimination task, significant correlations were observed from 70 to 840 ms (Figure 8A). In the texture discrimination task, three distinct clusters were identified (70-450 ms, 480-600 ms, and 630-850 ms; Figure 8B). Notably, direct task comparison revealed a marginally significant enhancement of V1–MEG correspondence in the 650-750 ms window (t(31) = 2.75, *q* = 0.0587, two-tailed, FDR corrected; Figure 8C).

**Figure 8:**
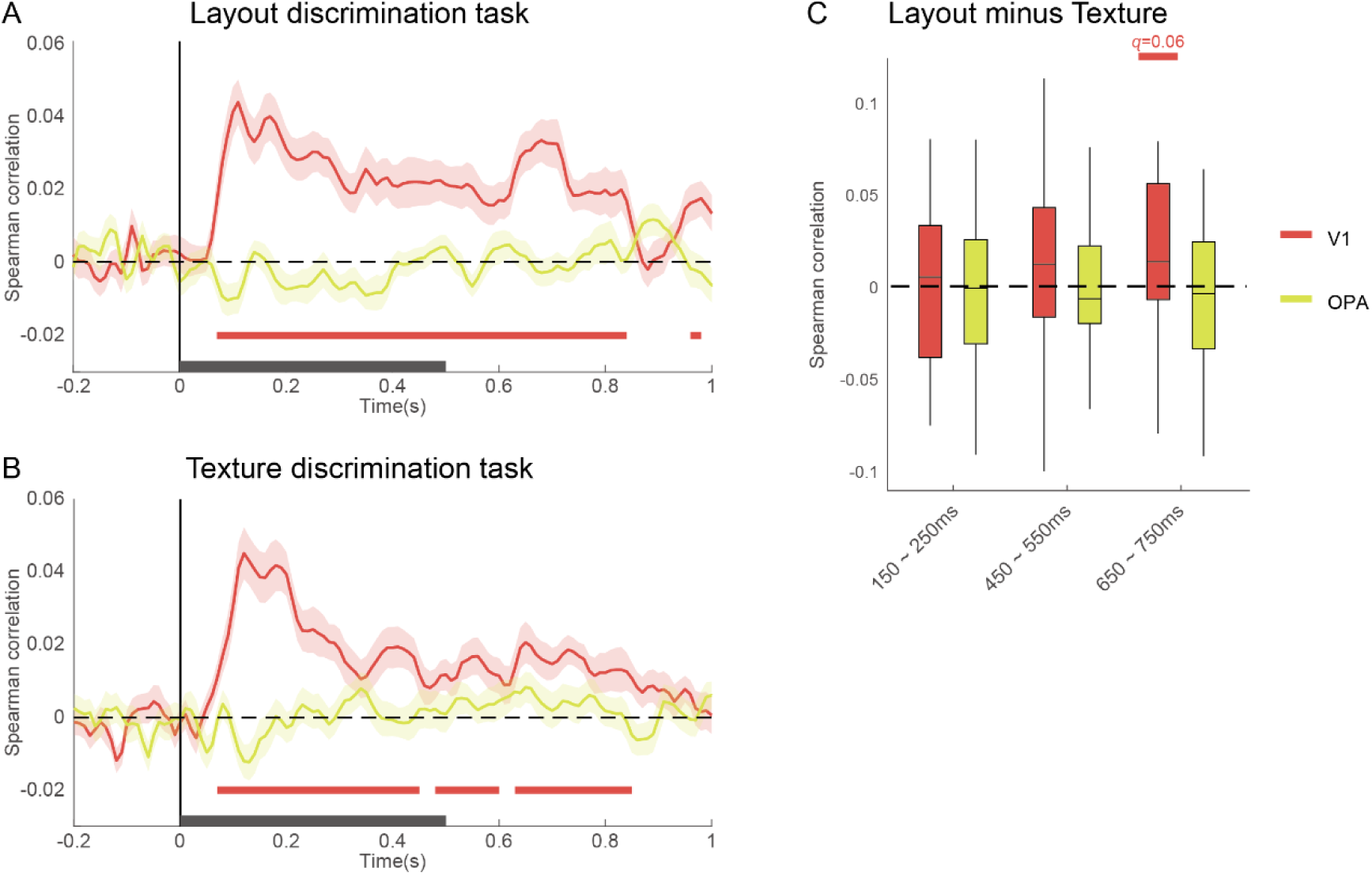
Cross-modal representational similarity between fMRI and MEG. Partial correlation between fMRI RDMs (V1, OPA) and the MEG RDMs from occipital channels for **A** layout discrimination task and **B** texture discrimination task. Significance was determined using one-sample t-tests against chance level, corrected for multiple comparisons with TFCE (one-tailed *p* < 0.05). Colored horizontal bars indicate significant clusters, and black bars on the x-axis denote image presentation period. **C**, Task-dependent changes in partial correlation across three critical time windows.

This late enhancement coincides temporally with the orientation-specific feedback effects observed in the MEG analyses (Figure 7C and 7D), providing convergent evidence for navigation-related feedback to early visual cortex during active layout processing.

## Discussion

The present study investigated how the human brain represents critical 3D layout features in naturalistic indoor scenes. We combined large-scale fMRI analysis, based on a new computer vision–based reconstruction approach, with targeted fMRI and MEG experiments using ground-truth annotated natural scenes. This multimodal approach enabled us to characterize fine-grained neural representations of boundary-related information at both the spatial and temporal levels.

Our findings provide converging evidence for a spatiotemporal dissociation between boundary distance and orientation representations. Early visual areas primarily encoded the relative distance to boundaries, whereas scene-selective areas were more engaged in representing boundary orientation. Integrating fMRI and MEG results further revealed distinct temporal profiles underlying these two forms of spatial coding: distance encoding emerged rapidly and remained task-invariant, peaking around 200 ms after image onset; orientation encoding appeared later, reaching its peak after 400 ms, and was strongly modulated by navigation-relevant task demands. Together, these results suggest a hierarchical organization of spatial layout representation along both cortical and temporal dimensions.

### Methodological advances

Previous studies on wall or boundary representation have faced two major methodological challenges that limit their ecological validity in the context of real-world navigation. First, most studies employed simplified, artificial stimuli consisting of isolated walls or geometric primitives. Although such stimuli facilitate precise experimental control, they lack critical real-world image regularities, such as semantic context, natural lighting conditions, and visual complexity (Groen & Baker, 2019). Conversely, naturalistic scene images provide more realistic inputs but lack quantitative ground-truth 3D layout information. Moreover, artificial scenes typically contain few or no objects, allowing for unobstructed visibility of wall contours. In contrast, naturalistic images include numerous objects that fragment the visibility of wall boundaries, making edge-based wall segmentation particularly challenging. To overcome these gaps, our study developed a computer vision-based approach that accurately estimates boundary distance and orientation from natural indoor scenes, enabling their direct application to large-scale datasets such as the NSD. With modest annotation effort, this approach enables efficient testing and comparing 3D layout models across existing public neuroimaging datasets, thereby bridging the methodological divide between controlled and naturalistic paradigms.

Second, many previous studies relied on tasks that bore limited relevance to navigation, potentially underestimating top-down modulations in spatial representation. Growing evidence indicates that scene-selective areas are strongly influenced by navigational context (Aminoff & Tarr, 2021; Bonner & Epstein, 2017; Chaisilprungraung & Park, 2021; J. Park & Park, 2020). For instance, Bonner & Epstein (2017) showed that OPA can distinguish doors from shape-compatible paintings, while Aminoff & Tarr (2021) demonstrated enhanced PPA and OPA activity when viewing scenes through windows versus picture frames. Our study directly builds upon these insights by incorporating tasks that vary in navigational relevance, thereby providing compelling evidence for task-dependent representation of 3D layout features.

### Implications in spatial layout processing

Inspired by the principle of vector coding (Bicanski & Burgess, 2020), our study modeled egocentric boundary features in terms of their distance and orientation relative to the observer. A key finding is the clear dissociation between early visual areas and scene-selective areas in representing these two layout components, suggesting a hierarchical process for extracting spatial layout information from visual input.

Early visual areas exhibited robust sensitivity to the relative distance, even after controlling for 2D visual and semantic features. The receptive fields in these areas are relatively small, making them well-suited for capturing local information. The fact that relative distance model, but not precise distance model (Supplementary Figure 4), captured the response pattern in these areas suggest that early visual cortex may play an essential role in estimating distance by segmenting the 2D image based on wall positions. This may explain the superior performance of our relative distance model in early visual areas compared with other alternatives (Supplementary Figure 2), as those models primarily focus on global features and lack egocentric directional information.

In contrast, orientation encoding likely involves more integrative computations, such as evaluating deviations in relative distances between adjacent image regions. Representing wall orientation provides an efficient means to infer navigational structure, since side walls in indoor environments are typically orthogonal and exhibit minimal curvature. Consistent with prior findings implicating scene-selective areas such as OPA in layout representation (Epstein & Kanwisher, 1998; Ferrara & Park, 2016; Henriksson et al., 2019; Julian et al., 2016; Kamps et al., 2016; Lescroart & Gallant, 2019; J. Park & Park, 2020), our results demonstrate that boundary orientation constitutes the key 3D layout feature encoded in these areas. Orientation reconstructing thus represents a crucial step in the transformation from 2D visual input to 3D structural understanding during navigation. Future studies should aim to elucidate the neural and computational mechanisms underlying this transformation.

Temporal dynamics from MEG data further reinforced this dissociation. Relative distance encoding emerged rapidly (∼200 ms), consistent with early feature extraction that is largely automatic and task-invariant, as shown in previous studies on natural scene perception (Greene & Hansen, 2018, 2020; Groen et al., 2016, 2017; Wischnewski & Peelen, 2021). Orientation encoding, however, manifested later, persisting between 400–700 ms, beyond the typical window for initial scene processing (Brandman & Peelen, 2023; Groen et al., 2016; Kaiser et al., 2019; Wu & Li, 2024). We propose that this later-stage processing reflects a refined representation of layout information, potentially through feedback mechanisms or working memory engagement, to support task-related spatial reasoning.

The observed task-dependent enhancement in both the Matterport3D fMRI and MEG experiments, centered around 700 ms, further supports this view. This effect, localized to V1 and posterior occipital regions, suggests feedback-mediated maintenance of detailed boundary information in early visual cortex, consistent with prior evidence that early visual areas support retention of sensory-like representation during visual working memory (Harrison & Tong, 2009; Jia et al., 2021; Kwak & Curtis, 2022; Pasternak & Greenlee, 2005; Rademaker et al., 2019; Serences et al., 2009) and visual imagery (Bainbridge et al., 2021; Dijkstra et al., 2018, 2019; Xie et al., 2020).

### Neural representation of self-pose

By leveraging recovered camera parameters from the NSD dataset, we found that pitch angle, but not roll angle, was robustly encoded in early visual and scene-selective areas. Pitch angle is a critical parameter for perceiving and navigating within 3D environments, influencing both spatial updating and memory for scenes (Wu & Li, 2024). The present findings provide the first direct evidence that pitch angle is explicitly encoded in the human visual cortex.

Despite comparable variability across images, roll angle was not reliably represented in any regions examined. This absence likely reflects ecological and behavioral constraints: roll head movements are rare and less relevant for everyday navigation, whereas pitch adjustments are essential for estimating elevation changes or assessing multi-level spaces. Consequently, the visual system may prioritize encoding pitch over roll due to its greater functional significance for spatial orientation.

### Absence of precise distance representation

Although scene-selective areas have been reported to encode distance and openness (Amit et al., 2012; Harel et al., 2013; Kravitz et al., 2011; Lescroart & Gallant, 2019; S. Park et al., 2011, 2015), we found no evidence for precise distance coding within our experimental design. This discrepancy likely reflects differences in spatial scale: most of our stimuli depicted near-range indoor scenes (1–10 m; Kravitz et al., 2011), whereas prior studies examined broader distance ranges across open environments (1–100 m). The visual system might adopt distinct strategies for encoding distance information across different spatial scales. Moreover, estimating precise distance requires at least one known coordinate in the world coordinate system, such as the real-world size of an object or the camera’s height. The brief static viewing (500 ms, or one saccade during natural movement) likely provides insufficient cues or processing time for such metric recovery.

Future research employing dynamic naturalistic video with depth annotations may help clarify this issue.

## Conclusions

In summary, we developed quantitative methods for reconstructing 3D layout information from naturalistic indoor scene and used it to reveal the neural architecture supporting human spatial layout perception. Our findings demonstrate that the human visual system represents boundary distance and orientation through a hierarchically organized process: engaging early visual areas for egocentric distance estimation and scene-selective regions for orientation encoding. These representations are further modulated by navigational relevance, highlighting the dynamic interplay between perception and action in the human brain during spatial navigation.

## Methods

### NSD experiment

#### The Natural Scenes Dataset

We used fMRI data from the Natural Scenes Dataset (NSD), which is described in detail by Allen et al. (2022). The dataset includes whole-brain fMRI measurements from 8 participants (6 females; age range: 19–32). Informed written consent was obtained from all participants, and the experimental protocol was approved by the University of Minnesota institutional review board. Over the course of a year, each participant viewed 9,000–10,000 colored natural scene images across 30–40 scan sessions. The full image set consists of 73,000 images, but only a subset of 1,000 images was viewed by all participants. These images were sourced from the Microsoft Common Objects in Context (COCO) database (Lin et al., 2014) and were presented for 3 seconds, following a 1-second gap between trials. Each image was shown three times, resulting in approximately 30,000 trials per participant. In each trial, participants performed a recognition task, determining whether the image had been presented in any previous session. The images were displayed at a visual angle of 8.4° × 8.4°, with participants instructed to focus on a central fixation dot superimposed on the image.

### Stimuli

#### Indoor NSD images

For our analysis, we selected indoor images from the NSD dataset using the COCO panoptic segmentation labels. These labels, designed for semantic segmentation in computer vision, delineate regions corresponding to both objects (thing) and background (stuff) within images. The panoptic labels consist of two main categories: "thing" (individual objects) and "stuff" (background elements like walls, floors, and ceilings). We focused on images that met the following criteria: (1) the image contained at least two of the three major layout components—ceiling, walls, and floor; (2) the walls were orthogonal to each other; and (3) no more than three side walls were present.

To ensure the walls were clearly visible in the selected images, we applied further manual screening to exclude images that met any of the following criteria: (1) blurred outlines of the background, (2) large objects or people obscuring the center of the image, or (3) incomplete enclosures (e.g., open or partially visible rooms).

#### Reconstructing camera’s internal and external parameters

The reconstruction of camera’s parameters is based on the computation of vanishing points (Orghidan et al., 2012). Six human annotators were recruited for the annotation task. Each annotator labeled a subset (mean = 292) of the filtered images and each image had one annotation. As shown in Figure 1A-(1), the annotators were instructed to label two pairs of lines in the selected images. Each pair of lines marks two clear edges that are parallel to the ground and to each other in the space shown in the image. The two pairs need to be orthogonal to each other. The line-pairs can be found at the edges of the walls, tiles on the floor, or any squared surface such as tables. The intersection of a line-pair is the vanishing point where the parallel lines converge in the 3D space. The straight line connecting the two vanishing points (*V*_1_ and *V*_2_) constitutes the horizon line *H* of the space (Supplementary Figure 1-(1)).

The camera’s internal and external parameters can help to map 2D image coordinates to 3D world coordinates with the equation:

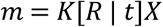

where *m* is the point in the camera coordinate system, *X* is the same point in the world coordinate system, *K* is the internal parameters and [*R* | *t*] is the external parameters: *R* is the rotation matrix and *t* is the translation matrix of the camera. The internal parameters contain focal length, skewness, and principal point in the form:

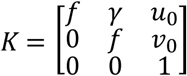

where *f* is the focal length, *γ* is the skewness, and point (*u*_0_,*v*_0_) is the location of principal point. The external parameters contain the camera pose, including yaw, pitch, roll, and translation of the camera. The external parameters can be written as a 3×4 transformation matrix.

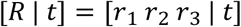

where *r*_1_, *r*_2_, *r*_3_ represent columns in the rotation matrix and *t* represents the translation. To simplify, the present study took the skewness, yaw, and translation to be zero and considered the principal point to be located at the center of the image. With the simplification, the focal length can be calculated as:

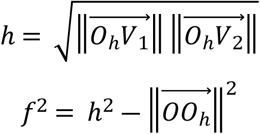

where *O*_ℎ_ is the projection point of the principal point to the horizon line *H* and *V*_1_, *V*_2_ are the two vanishing points (Figure 1A-(2) and Supplementary Figure 1-(2a)).

Since we restricted the yaw angle to be zero, the pitch and roll angles can be calculated by the rotation matrix determined by the vanishing point *V*_*s*_ where the parallel lines extending straight front converge. *V*_*s*_ can be located as the intersection of the horizon line *H* and the vertical line through the principal point *O*. The paired vanishing point *V*_*c*_ of the direction of left and right can be easily located with the equation (Supplementary Figure 1-(2b)):

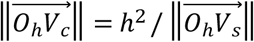

Given the locations of *V*_*s*_ and *V*_*c*_ and the focal length, the external parameters can be calculated as:

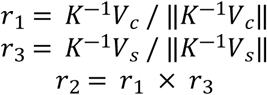

and the pitch and roll angles are

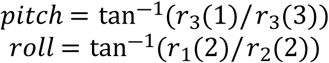

#### Reconstructing scene layout

Another human annotator was recruited for the annotation task and each image had one annotation. As shown in Figure 1A-(4), the annotator was told to label two lines based on the number of side walls present: (1) If the image contains three side walls, label the two lines on the middle side wall. (2) If there are fewer than three side walls, label the two lines on any side wall.. The first line *L_1_* marks an edge of the wall that is parallel to the ground. The two endpoints of the first line need to be located at the corners between the annotated wall and the side walls. The second line *L_2_* marks the height of the annotated wall. The endpoints of the second line need to be located at the intersection lines between the annotated wall and the ceiling and the floor, respectively.

The reconstruction of scene layout needs to meet the prerequisites that all walls are orthogonal to each other and there are at most three side walls. Thus, all images that do not meet this prerequisite were discarded. In brief, the reconstruction includes the following steps: locating the third vanishing point, outlining the annotated wall, and outlining other side walls.

**Locating the third vanishing point.** The third vanishing point *V*_3_ is the converging point of the parallel lines that are orthogonal to the ground. The principal point *O* is the orthocenter of the triangle that is formed by *V*_1_, *V*_2_ and *V*_3_. Thus, the third vanishing point *V*_3_ can be easily calculated (Figure 1A-(3) and Supplementary Figure 1-(3)).

**Outlining the annotated wall.** According to the prerequisite, the conjunctions between the annotated wall and the ceiling and floor are parallel to the ground, and the conjunctions of the annotated wall and other side walls are orthogonal to the ground. The conjunctions of the annotated wall and the ceiling and floor intersect at the intersection point of the first line *L_1_* and the horizon line *H*. The conjunctions of the annotated wall and other side walls intersect at *V*_3_. The detailed steps are following:

1. Locate the intersection point *V*_*a*_ of *L_1_* and *H*.
2. Connect *V*_*a*_ to the two endpoints of the second line *L_2_*. These lines are the conjunctions of the annotated wall and the ceiling and floor.
3. Connect *V*_3_ to the two endpoints of the first line *L_1_*. These lines are the conjunctions of the annotated wall and other side walls. The intersection points of these two lines and the ceiling are named as *p*_2_ and *p*_3_. The intersection points of the two lines and the floor are named as *p*_6_ and *p*_7_.

The quadrilateral decided by *p*_2_, *p*_3_, *p*_6_ and *p*_7_ is the outline of the annotated wall (Figure 1A-(4)).

**Outlining other side walls.** Other side walls are orthogonal to the annotated wall. The intersection point *V*_*o*_ of other walls and *H* follows the equation:

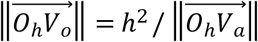

and *V*_*o*_ thus can be easily located. We outlined other side walls with the following steps:

1. Locate the intersection point *V*_*o*_ using the equation.
2. Connect *V*_*o*_ to *p*_2_ and *p*_6_. These lines are the conjunctions of the left wall and the ceiling and floor. The intersection points of these two lines and the edges of the image are named as *p*_1_ and *p*_5_. If *p*_2_ and *p*_6_ are on the edges of the image, skip this step.
3. Connect *V*_*o*_ to *p*_3_ and *p*_7_. These lines are the conjunctions of the right wall and the ceiling and floor. The intersection points of these two lines and the edges of the image are named as *p*_4_ and *p*_8_. If *p*_3_ and *p*_7_ are on the edges of the image, skip this step.

The area to the left of the polyline decided by *p*_1_, *p*_2_, *p*_5_ and *p*_6_ is the left wall. The area to the right of the polyline decided by *p*_3_, *p*_4_, *p*_7_ and *p*_8_ is the right wall (Figure 1A-(5,6)).

**Orientation of side walls.** The orientation angle relative to the front is decided by the point *V*_*a*_ and *V*_*o*_.

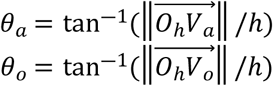

where *θ*_*a*_ is the orientation angle of annotated wall and *θ*_*o*_ of orthogonal walls.

### Feature spaces

#### GIST

The GIST feature quantifies spatial frequencies and orientations across different locations in a scene (Oliva & Torralba, 2001). For each scene image, we first converted it to grayscale and then processed it using 512 filters (or channels), which consisted of 4 spatial frequencies and 8 orientations at 16 spatial locations (4 rows × 4 columns). To reduce the dimensionality of the resulting data, we performed principal component analysis (PCA) on all 73,000 NSD images. The top 59 principal components accounted for over 95% of the variance and were retained as the GIST feature space for each image.

#### Texture

To quantify the mid-level features of scene images, we used the texture model proposed by Portilla & Simoncelli (2000), which has been shown to predict both human perceptual judgments and neural responses to textures (Freeman & Simoncelli, 2011; Long et al., 2018). First, each scene image was converted to grayscale and resampled to a resolution of 256 × 256 pixels. The images were then decomposed into orientation and spatial frequency sub-bands using the steerable pyramid transformation. These sub-bands represent spatial maps of complex-valued coefficients, where the real part corresponds to the response of V1 simple cells and the magnitude of the coefficients corresponds to the response of V1 complex cells.

The model computed several descriptors within and across these sub-bands to capture texture features. Specifically, the descriptors included: (1) pixel autocorrelations at different positions within each sub-band, which characterize periodicity; (2) magnitude autocorrelations and cross-correlations across sub-bands, which capture structural elements like edges and corners, as well as second-order texture properties; and (3) cross-correlations of the real part of the sub-bands with phase-doubled responses at the next coarser scale, which distinguish edges from lines and capture gradients of shading.

Each scene image was processed through the texture model using 4 orientations and 4 spatial frequencies across a 5 × 5 neighborhood at each of the 4 (2 rows × 2 columns) spatial locations. We performed principal component analysis (PCA) on the texture parameters derived from all 73,000 NSD images. The top 125 principal components accounted for over 95% of the variance and were retained as the texture feature space for each image.

#### Semantic

To model the semantic properties of the scene images, we employed the object2vec model (Bonner & Epstein, 2021), which is based on word2vec, a well-established algorithm in computational linguistics used to model the lexical-semantic properties of words based on their co-occurrence statistics (Mikolov et al., 2013). Object2vec extends this idea to image semantics by learning to represent image labels in a continuous vector space. Specifically, the method uses the continuous bag of words (CBOW) model of word2vec, where the target label is predicted based on the surrounding context labels within an image.

In this study, we trained the object2vec model on the training set of the COCO database, which contains 118,287 images. For training, each image was converted into a "sentence" consisting of its panoptic segmentation labels. The panoptic segmentation labels in COCO are divided into 80 "thing" categories (e.g., objects such as people, cars) and 53 "stuff" categories (e.g., background regions like walls, sky), totaling 133 labels.

The object2vec embeddings were learned by training the CBOW model of word2vec with an embedding dimension of 50. The model was trained for 100 epochs, using a negative sampling parameter of 20 and a window size of 15, meaning that the model would consider 30 surrounding labels within each image to predict the target label. After training, the object2vec model generated a 50-dimensional vector to represent each label. The semantic feature space for each image was obtained by averaging the vectors of all its labels, providing a compact representation of its semantic content.

#### Layout orientation model

The layout orientation model represents the orientations of the side walls (i.e., the boundaries) in a scene (Figure 2C left). Using the aerial view annotations, we calculated the orientations and start and end points of the side walls. For each side wall, we computed its normal vector—unit vector that is perpendicular to the wall and directed towards the camera. These normal vectors are represented in a right-handed 2D coordinate system, with the +x axis pointing to the right and the +y axis pointing forward.

To discretize the orientation information across different locations, we quantized the normal vectors into 5 histogram bins. These bins are organized along the horizontal axis in the perspective of observer, evenly dividing the entire field of view. The orientation information of the side walls is integrated across these location bins using the following equations:

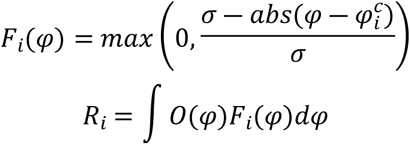

where *σ* is the angular width of each bin, *φ*^*c*^is the center of the *i*-th bin, and *φ* represents any direction in the field of view. The responses of the bins are computed by integrating over the entire field of view, where *F*_*i*_(*φ*) is the soft-histogram function for the *i*-th bin, and *O*(*φ*) corresponds to the orientation angle of the wall at the direction *φ*.

#### Layout relative distance model

Assuming that all walls are orthogonal to each other, the proportion of the side wall relative to the entire pixel space provides a rough estimate of the relative distance of the side walls. As the proportion of side wall pixels increases, the perceived distance to the side walls decreases.

In the relative distance model (Figure 2C right), each scene image is divided into 5 equal horizontal bins. For each bin, the proportion of side wall pixels is calculated based on the layout segmentation map in the pixel space. This proportion serves as a coarse measure of the relative distance of the side walls within each bin.

### MRI data acquisition and preprocessing

Functional data were acquired at 7T using whole-brain gradient-echo echo-planar imaging (EPI) with an isotropic resolution of 1.8 mm and a repetition time (TR) of 1.6 s. The functional data were preprocessed by performing temporal interpolation to correct for slice timing and spatial interpolation for head motion correction. A general linear model (GLM) approach was applied to estimate single-trial beta weights. The third beta version (betas_fithrf_GLMdenoise_RR) was used in this study. In brief, a new GLMsingle algorithm (Prince et al., 2022) was used to estimate voxel responses. This algorithm implements optimized denoising and regularization procedures to accurately measure changes in brain activity evoked by experimental stimuli. For further details on this methodology, please refer to Allen et al. (2022).

### Defining ROIs

Retinotopic and category-selective regions of interest (ROIs) were defined based on functional localizers from the NSD experiment. The category localizer task was employed to identify scene-selective areas, including the parahippocampal place area (PPA), occipital place area (OPA), and retrosplenial cortex (RSC), as well as the face-selective fusiform face area (FFA). A pRF (population receptive field) mapping task was used to define the retinotopic visual area V1.

### Decoding analysis

For each participant, a support vector regression (SVR) model was trained to decode the neural representation of pitch and roll angles. SVR is a machine learning algorithm commonly used for regression analysis and is particularly effective in decoding continuous variables (e.g., Schubert et al., 2021). The epsilon SVR was implemented using LibSVM with a linear kernel. A ten-fold cross-validation procedure was employed to assess the performance of the SVR model. Decoding performance was quantified by computing the Pearson correlation between the predicted and actual angles. The chance level was defined as zero, indicating no correlation between the predicted and actual angles. Group-level statistical analysis was conducted using a one-tailed t-test against the chance level. False discovery rate (FDR) correction was applied to adjust for multiple comparisons across ROIs.

### Representational similarity analysis

Neural representational dissimilarity matrices (RDMs) were constructed within each ROI for each participant. Each scene image was presented up to three times. Multivoxel activation patterns for each image were obtained by averaging all trials of the image. The distances between resulting patterns were calculated using Pearson correlation, which was then used to construct the RDMs.

Model RDMs were generated for all the feature spaces described previously. The RDMs of GIST and texture features were calculated using Euclidean distance, while the RDM of semantic features was measured using cosine distance. The RDMs of the layout orientation and relative distance models were measured using city block distance.

To identify the neural representation of the feature spaces, we compared the neural and model RDMs for each participant. A partial correlation analysis was applied to characterize the coding of layout, while controlling for perceptual and semantic scene features. We incorporated the models of layout orientation and relative distance, while controlling for GIST, texture, and semantic features. One-tailed t-tests were used across participants to assess the partial effects for statistical significance, with FDR correction applied for multiple comparisons across all models and ROIs.

### Searchlight analysis

The searchlight analysis was conducted in the native space of each participant with the mask comprised of ‘nsdgeneral’, ‘floc-places’, ‘floc-faces’, and ‘floc-bodies’ provided in the NSD dataset (Allen et al., 2022). The mask covered voxels responsive to the NSD experiment in the posterior aspect of cortex and fROIs in the native space. We moved a cubic searchlight in the radius of 5 mm in the mask. At each voxel, we obtained the partial correlations of orientation and relative distance models while controlling for image features. The searchlight results were transformed to surface space for visualization using the interpolation of nearest-neighbor.

## Matterport3D fMRI experiment

### Participants

Thirty healthy adults (mean age = 21.53 years, SD = 2.16; 11 females) with normal or corrected-to-normal vision participated in the study. All participants provided written informed consent before the experiment and were compensated for their time. The study was approved by the Committee for Protecting Human and Animal Subjects at the School of Psychological and Cognitive Sciences at Peking University (Institutional Review Board Protocol No: 2023-08-03).

### Stimuli

#### Naturalistic scene images

We selected 60 naturalistic indoor images from the Matterport3D database, a large-scale RGB-D dataset comprising 10,800 panoramic views generated from 194,400 RGB-D images across 90 building-scale environments (Chang et al., 2017). The database provides depth map, camera pose, and surface normal map for each image. Additionally, ground-truth layout annotations were sourced from the Matterport3D-Layout dataset, a specialized database for scene layout estimation (Zhang et al., 2020).

The images were categorized into five indoor scene types: bathroom, bedroom, kitchen, living room, and hallway. To maintain consistency, all selected images had a pitch and roll angle of 0° and were extracted from panoramic views with a horizontal field of view (FOV) of 60° and a vertical FOV of 53.3°. The final images were resized to 650 × 520 pixels for experimental use.

#### Synthesis of layout segmentation map

We generated a ‘ground-truth’ segmentation map along with two additional segmentation maps, each incorporating a 10° yaw angle deviation to the left and right relative to the original image.

The process began by reconstructing an aerial view map of the original image using the depth and normal maps provided by the Matterport3D database. Based on this reconstructed aerial view, we generated aerial view maps with yaw angle deviations of ±10°. Given these aerial view maps and the ceiling height information from the Matterport3D database, the segmentation maps were synthesized using the method described in the Reconstructing Scene Layout section of the NSD experiment.

Consistent with the labeling approach in the NSD experiment, we assigned labels to the side walls in the layout segmentation maps. The side wall regions were labeled sequentially from left to right.

#### Synthesis of texture image

For each image, we synthesized both an original texture image and a feature-swapped texture image.

The original texture images were generated to match a set of descriptors derived from the texture model used in the NSD experiment for the original image. Specifically, the model’s responses to the original image were computed using four orientations and four spatial frequencies across a 7 × 7 neighborhood at each of the nine spatial locations (3 rows × 3 columns). An image of Gaussian white noise was then iteratively adjusted to match these model responses. This synthesis procedure approximates sampling from the maximum entropy distribution of images that conform to a given set of model responses. To generate the texture images, we applied gradient descent for 50 iterations.

To synthesize the feature-swapped texture image, we replaced the pixel autocorrelation descriptor of the original image with that of a randomly selected image from the Matterport3D database, ensuring that this selected image had not been presented in the task. The same iterative procedure was then applied to match the swapped model responses.

### Feature spaces

We employed the GIST, texture, and semantic models to quantify image features. The GIST and texture features were computed as the NSD experiment using same model parameters. The responses of texture models were estimated among all scene images with a pitch angle of 0° in the Matterport3D database, in total of 64,800 scene images. Then we performed the PCA on the responses of texture models. The top 118 principal components accounted for over 95% of the variance and were retained as the texture feature space for each image. The responses of GIST model were directly used without PCA. The semantic feature was computed by the panoptic labels rendered from the ‘house segmentations’ in the Matterport3D database using the camera parameters. An object2vec model was performed as the NSD experiment using the same model parameters except for an embedding dimension of 20. For layout representation, we incorporated the two layout models described in the NSD experiment. The relative distance model was computed from the ground-truth layout segmentation map in the Matterport3D-Layout dataset and the orientation model was computed from the aerial view map, reconstructed using the segmentation, depth, and normal maps.

### Experimental design

All experiments were conducted using Psychtoolbox software. Stimuli were displayed on an LCD monitor (refresh rate, 60 Hz; resolution, 1920 ×1080), subtending a visual angle of 9.5° × 7.6°. Participants viewed the stimuli at a distance of 182 cm through a mirror placed above their eyes. Each participant completed two scanning sessions, with each session consisting of three runs of the layout discrimination task and three runs of the texture discrimination task. Additionally, two localizer runs were performed before task runs in the first session.

#### Layout discrimination and texture discrimination tasks

We employed a match-to-sample paradigm for both tasks (Figure 4A). Each trial commenced with a fixation cross displayed for 1–3 seconds, followed by a scene image presented for 0.5 seconds. After an additional fixation period of 3–5 seconds, the response phase began, during which two stimuli appeared on the left and right sides of the screen. Participants were instructed to select the stimulus that best matched the initial scene image by pressing a button using their right hand. The response window lasted 3.5 seconds; however, the stimuli disappeared once a response was made. Inter-trial intervals were jittered and pseudo-randomized. To ensure adequate baseline signal, two 16-second rest periods were included at the beginning and end of each run. Each run comprised 40 trials, totaling 482 seconds in duration.

In the layout discrimination task, participants were presented with two layout segmentation maps during the response phase: one representing the ’ground-truth’ layout of the scene and another with a deviation. Participants were instructed to choose the map that accurately depicted the scene’s layout. In the texture discrimination task, participants were shown two texture images: one corresponding to the original scene and another feature-swapped version. They were asked to select the image that best matched the scene’s color distribution.

Within each session, participants completed all 3 runs of one task before proceeding to the next task, with the order of tasks counterbalanced across sessions. Each scene image was presented twice per task in a session, ensuring no image was repeated within a single run.

#### Functional localizer runs

We employed a block design to define the functional regions of interest (fROIs) for each participant. The localizer stimuli consisted of colored images from four categories: scenes, faces, objects, and scrambled objects. Images within the same category were presented in centrally displayed blocks lasting 16 seconds. Each block contained 20 images, with each image presented for 300 ms, followed by a 500 ms blank interval. Inter-block intervals lasted 8 seconds.

Participants performed a one-back task, in which they were required to press a button when two consecutive images were identical. Each localizer run lasted 408 seconds and comprised four blocks per category. The order of category blocks was counterbalanced across participants and runs.

### MRI data acquisition

BOLD fMRI data were collected using a 3T Siemens Prisma scanner equipped with a 64-channel receiver head coil. Functional images were acquired using a multiband echo-planar imaging (EPI) sequence with the following parameters: multiband factor = 2, repetition time (TR) = 2 s, echo time (TE) = 30 ms, matrix size = 112 × 112 × 62, flip angle = 90°, spatial resolution = 2 × 2 × 2.3 mm³, and number of slices = 62.

Before the functional scans in each session, a high-resolution T1-weighted 3D anatomical dataset was acquired for each participant to aid in image registration. The anatomical scans were obtained using an MPRAGE sequence with the following parameters: TR = 2530 ms, TE = 2.98 ms, matrix size = 448 × 512 × 192, flip angle = 7°, spatial resolution = 0.5 × 0.5 × 1 mm³, number of slices = 192, and slice thickness = 1 mm.

### fMRI pre-processing

The anatomical and functional data were pre-processed and analyzed using AFNI (Cox, 1996). Functional images were slice-time corrected using the AFNI function 3dTshift and motion-corrected to the reference image, which was considered the minimum outlier (using 3dVolreg). The functional images from each session were aligned with the corresponding anatomical images. The functional data from the second session were then co-registered and resampled to match the space of the first session, guided by the alignment of the two anatomical images. After co-registration, the signal amplitudes of the functional images were rescaled to a 0-200 range. All RSA analyses were conducted in the original space.

The responses from the task runs were modeled using a General Linear Model (GLM). Each trial was modeled with a canonical Hemodynamic Response Function (HRF), derived from the onset of the original image, and convolved with a 0.5-second square wave. Additional regressors included the participant’s response, six motion parameters, and three polynomial terms to account for slow signal drifts. We performed the least-squares-sum (LSS) estimation within AFNI to obtain single-trial beta estimates (Mumford et al., 2011). These single-trial beta values were then normalized within each run for subsequent analyses.

### Defining ROIs

Using the functional localizer runs, we defined three scene-selective regions of interest (ROIs) in each hemisphere: the parahippocampal place area (PPA), the occipital place area (OPA), and the retrosplenial complex (RSC). We fitted the response model using a General Linear Model (GLM) in AFNI (3dDeconvolve and 3dREMLfit). BOLD responses were modeled by convolving a standard Hemodynamic Response Function (HRF) with a 16-second square wave for each category. Estimated motion parameters, participants’ responses, and three polynomial terms to account for slow drifts were included as regressors of no interest.

Scene-selective areas were defined as contiguous clusters of voxels with a threshold of p < 1×10⁻⁴ (uncorrected) under the contrast of scene > face, based on their anatomical locations. Specifically, the PPA was defined by locating the cluster between the posterior parahippocampal gyrus and the lingual gyrus, the OPA was defined near the transverse occipital sulcus, and the RSC was located near the posterior cingulate cortex. In seven participants, the localizer failed to identify the RSC. The primary visual cortex (V1) was defined based on each participant’s anatomical parcellation from Freesurfer (Hinds et al., 2009).

### Representational similarity analysis

For each participant, neural representational dissimilarity matrices (RDMs) were constructed for each task within each region of interest (ROI). Each image was presented four times per task. Multivoxel activation patterns for each image were derived by averaging the trials across sessions. The distances between patterns were calculated as one minus the Pearson correlation.

Model RDMs were constructed for the image feature models, layout orientation model, and layout relative distance model using the same distance measure as in the NSD experiment.

The partial correlation analysis, similar to those in the NSD experiment, was performed separately for the two tasks to characterize the task-dependent representations of layout. We included the models of layout orientation and layout relative distance in the analysis, again controlling for the three image feature models. One-tailed t-tests were applied across participants to assess the statistical significance of partial effects, with false discovery rate (FDR) correction for multiple comparisons across all models and ROIs.

### Matterport3D MEG experiment

#### Participants

Thirty-two healthy adults (mean age = 21.22 years, SD = 2.51; 20 females) with normal or corrected-to-normal vision participated in the study. All participants provided written informed consent before the experiment and were compensated for their time. The study was approved by the Committee for Protecting Human and Animal Subjects at the School of Psychological and Cognitive Sciences at Peking University (Institutional Review Board Protocol No: 2024-10-02).

#### Stimuli

The same set of 60 indoor scene images, along with their synthesized segmentation maps and texture images, were used in the MEG experiment as in the fMRI experiment. The stimuli were displayed on a rear-projection screen (refresh rate: 60 Hz) with a visual angle of 9.5° × 7.6°. Participants viewed the stimuli at a distance of 91 cm.

#### Experimental design

The design of the MEG experiment closely followed the match-to-sample paradigm from the fMRI experiment. Each trial began with a fixation period of 1-3 seconds, followed by the presentation of a scene image for 0.5 seconds. After another fixation period lasting 3-5 seconds, the response phase began, during which two stimuli were presented side-by-side on the screen. Participants were instructed to select the stimulus that best described the previously shown image within 3 seconds using their right hand.

Both the layout discrimination task and the texture discrimination task consisted of three runs each. All 60 scene images were presented in each run, with each image shown three times per task. All tasks were completed within a single session.

#### Data acquisition and preprocessing

Electromagnetic brain activity was recorded using an Eekta Neuromag 306 MEG system, which consists of 204 planar gradiometers and 102 magnetometers. The system includes 102 triple-sensor elements, with each sensor comprising one magnetometer and two planar gradiometers. Data were sampled continuously at 1000 Hz and band-pass filtered online between 0.1 and 330 Hz.

Offline preprocessing was performed using the MNE-Python package. Temporal signal space separation (tSSS) was applied to reduce environmental and head motion artifacts. Independent component analysis (ICA) was then used on the data with the fastICA algorithm implemented in MNE-Python. Components associated with eye blinks and saccades were identified and removed from the raw unfiltered data. Subsequently, the data were demeaned, detrended, and down-sampled to 100 Hz. A time window of 1700 ms was applied to segment the raw data, spanning from 200 ms before the first image onset to 1500 ms after the onset. Trials were band-pass filtered between 0.1 and 30 Hz and then excluded if the gradiometer value exceeded 5000 fT/cm or if the magnetometer value exceeded 5000 fT.

#### Representational similarity analysis

RSA in Matterport3D MEG experiment was conducted using the CoSMoMVPA toolbox. The event-related RSA was performed across 24 occipital magnetometers and one of their combined gradiometers. The selection of these channels was based on the extensive involvement of occipital regions in encoding layout, as observed in the previous two fMRI experiments. Temporal smoothing was applied by averaging over two adjacent time points (±20 ms). Neural RDMs were constructed at each time point for each task and participant. For each image, all trials were averaged, and distances between images were calculated as one minus the Pearson correlation coefficient.

The partial correlation analysis conducted was performed across all time points within the trial. The analysis incorporated the model RDMs of three image features, layout orientation, and layout relative distance.

To compare the correspondence between the representations in fMRI and MEG, we computed group-level fMRI-RDMs by averaging the RDMs of all participants for V1 and three scene-selective areas, respectively. Partial correlation analysis was then applied to compare the fMRI-RDMs of V1 and OPA with the MEG-RDMs of each participant at each time point.

Statistical significance was assessed against chance levels by computing random-effect temporal-cluster statistics, corrected for multiple comparisons. The null distribution was generated through t-tests over 10,000 iterations, in which the sign of decoding performance above chance was randomly flipped. Threshold-free cluster enhancement (TFCE) was used as the cluster statistic (Smith & Nichols, 2009), with a threshold step of 0.1. Significant temporal clusters were identified using a cluster-forming threshold of p < 0.05, one-tailed.

Time window-based RSA was performed across occipital magnetometers and one of their combined gradiometers. Here, we chose three 100-ms time windows corresponding to the significant clusters of layout models, centered at 200 ms, 500 ms, and 700 ms after image onset. All time points within each time window were averaged for each channel. The neural RDMs were constructed at each time window for each task and participant. The partial correlation analyses of model RDMs and fMRI-RDMs were conducted as the event-related RSA.

#### Searchlight analysis

The time window-based RSA method described above was applied in a searchlight approach across the entire scalp of each participant. All magnetometers and one of the combined gradiometers were involved in the searchlight analysis, resulting in 204 channels. For each channel, all time points within each time window were averaged. For each time window, task, and participant, the searchlight analysis was performed with a channel neighborhood determined by Delaunay triangulation. On average each location had a neighborhood of 16 channels. The same statistic test and TFCE procedures as described in the event-related RSA were applied to the partial correlation results across the scalp.

## Data availability

Data and script can be found at OSF at https://osf.io/uxwr4/

## Declaration of Interests

None declared.

## Acknowledgments

This study was supported by grants from STI2030-Major Projects (2021ZD0200204) and the National Natural Science Foundation of China (32271104).

## Notes

### Competing Interest Statement

The authors have declared no competing interest.

## Reference

1. Alexander, A. S., Carstensen, L. C., Hinman, J. R., Raudies, F., Chapman, G. W., & Hasselmo, M. E. (2020). Egocentric boundary vector tuning of the retrosplenial cortex. Science Advances, 6(8), eaaz2322. 10.1126/sciadv.aaz2322

2. Allen, E. J., St-Yves, G., Wu, Y., Breedlove, J. L., Prince, J. S., Dowdle, L. T., Nau, M., Caron, B., Pestilli, F., Charest, I., Hutchinson, J. B., Naselaris, T., & Kay, K. (2022). A massive 7T fMRI dataset to bridge cognitive neuroscience and artificial intelligence. Nature Neuroscience, 25(1), 116–126. 10.1038/s41593-021-00962-x

3. Aminoff, E. M., & Tarr, M. J. (2021). Functional Context Affects Scene Processing. Journal of Cognitive Neuroscience, 33(5), 933–945. 10.1162/jocn_a_01694

4. Amit, E., Mehoudar, E., Trope, Y., & Yovel, G. (2012). Do object-category selective regions in the ventral visual stream represent perceived distance information? Brain and Cognition, 80(2), 201–213. 10.1016/j.bandc.2012.06.006

5. Bainbridge, W. A., Hall, E. H., & Baker, C. I. (2021). Distinct Representational Structure and Localization for Visual Encoding and Recall during Visual Imagery. Cerebral Cortex, 31(4), 1898–1913. 10.1093/cercor/bhaa329

6. Barry, C., Lever, C., Hayman, R., Hartley, T., Burton, S., O’Keefe, J., Jeffery, K., & Burgess, N. (2006). The boundary vector cell model of place cell firing and spatial memory. Reviews in the Neurosciences, 17(1–2), 71–98.

7. Bicanski, A., & Burgess, N. (2020). Neuronal vector coding in spatial cognition. Nature Reviews Neuroscience, 21(9), Article 9. 10.1038/s41583-020-0336-9

8. Bonner, M. F., & Epstein, R. A. (2017). Coding of navigational affordances in the human visual system. Proceedings of the National Academy of Sciences, 114(18), 4793–4798. 10.1073/pnas.1618228114

9. Bonner, M. F., & Epstein, R. A. (2021). Object representations in the human brain reflect the co-occurrence statistics of vision and language. Nature Communications, 12(1), 4081. 10.1038/s41467-021-24368-2

10. Brandman, T., & Peelen, M. V. (2023). Objects sharpen visual scene representations: Evidence from MEG decoding. Cerebral Cortex, 33(16), 9524–9531. 10.1093/cercor/bhad222

11. Chaisilprungraung, T., & Park, S. (2021). “Scene” from inside: The representation of Observer’s space in high-level visual cortex. Neuropsychologia, 161, 108010. 10.1016/j.neuropsychologia.2021.108010

12. Chang, A., Dai, A., Funkhouser, T., Halber, M., Nießner, M., Savva, M., Song, S., Zeng, A., & Zhang, Y. (2017). *Matterport3D: Learning from RGB-D Data in Indoor Environments* (No. arXiv:1709.06158). arXiv. 10.48550/arXiv.1709.06158

13. Cox, R. W. (1996). AFNI: software for analysis and visualization of functional magnetic resonance neuroimages. Computers and Biomedical Research, 29(3), 162–173. 10.1006/cbmr.1996.0014

14. Dijkstra, N., Bosch, S. E., & van Gerven, M. A. J. (2019). Shared Neural Mechanisms of Visual Perception and Imagery. Trends in Cognitive Sciences, 23(5), 423–434. 10.1016/j.tics.2019.02.004

15. Dijkstra, N., Mostert, P., Lange, F. P. de, Bosch, S., & van Gerven, M. A. (2018). Differential temporal dynamics during visual imagery and perception. eLife, 7, e33904. 10.7554/eLife.33904

16. Epstein, R. A., & Baker, C. I. (2019). Scene Perception in the Human Brain. Annual Review of Vision Science, 5(1), 373–397. 10.1146/annurev-vision-091718-014809

17. Epstein, R. A., & Kanwisher, N. (1998). A cortical representation of the local visual environment. Nature, 392(6676), 598–601. 10.1038/33402

18. Ferrara, K., & Park, S. (2016). Neural representation of scene boundaries. Neuropsychologia, 89, 180–190. 10.1016/j.neuropsychologia.2016.05.012

19. Freeman, J., & Simoncelli, E. P. (2011). Metamers of the ventral stream. Nature Neuroscience, 14(9), Article 9. 10.1038/nn.2889

20. Greene, M. R., & Hansen, B. C. (2018). Shared spatiotemporal category representations in biological and artificial deep neural networks. PLOS Computational Biology, 14(7), e1006327. 10.1371/journal.pcbi.1006327

21. Greene, M. R., & Hansen, B. C. (2020). Disentangling the Independent Contributions of Visual and Conceptual Features to the Spatiotemporal Dynamics of Scene Categorization. The Journal of Neuroscience, 40(27), 5283–5299. 10.1523/JNEUROSCI.2088-19.2020

22. Groen, I. I. A., & Baker, C. I. (2019). Scenes in the Human Brain: Comparing 2D versus 3D Representations. Neuron, 101(1), 8–10. 10.1016/j.neuron.2018.12.014

23. Groen, I. I. A., Ghebreab, S., Lamme, V. A. F., & Scholte, H. S. (2016). The time course of natural scene perception with reduced attention. Journal of Neurophysiology, 115(2), 931–946. 10.1152/jn.00896.2015

24. Groen, I. I. A., Silson, E. H., & Baker, C. I. (2017). Contributions of low- and high-level properties to neural processing of visual scenes in the human brain. Philosophical Transactions of the Royal Society B: Biological Sciences, 372(1714), 20160102. 10.1098/rstb.2016.0102

25. Hafting, T., Fyhn, M., Molden, S., Moser, M.-B., & Moser, E. I. (2005). Microstructure of a spatial map in the entorhinal cortex. Nature, 436(7052), Article 7052. 10.1038/nature03721

26. Harel, A., Kravitz, D. J., & Baker, C. I. (2013). Deconstructing visual scenes in cortex: Gradients of object and spatial layout information. Cerebral Cortex, 23(4), 947–957. 10.1093/cercor/bhs091

27. Harrison, S. A., & Tong, F. (2009). Decoding reveals the contents of visual working memory in early visual areas. Nature, 458(7238), 632–635.

28. Henriksson, L., Mur, M., & Kriegeskorte, N. (2019). Rapid Invariant Encoding of Scene Layout in Human OPA. Neuron, 103(1), 161–171.e3. 10.1016/j.neuron.2019.04.014

29. Hinds, O., Polimeni, J. R., Rajendran, N., Balasubramanian, M., Amunts, K., Zilles, K., Schwartz, E. L., Fischl, B., & Triantafyllou, C. (2009). Locating the functional and anatomical boundaries of human primary visual cortex. Neuroimage, 46(4), 915–922. 10.1016/j.neuroimage.2009.03.036

30. Hinman, J. R., Chapman, G. W., & Hasselmo, M. E. (2019). Neuronal representation of environmental boundaries in egocentric coordinates. Nature Communications, 10(1), Article 1. 10.1038/s41467-019-10722-y

31. Jia, K., Li, Y., Gong, M., Huang, H., Wang, Y., & Li, S. (2021). Perceptual learning beyond perception: Mnemonic representation in early visual cortex and intraparietal sulcus. Journal of Neuroscience, 41(20), 4476–4486.

32. Julian, J. B., Ryan, J., Hamilton, R. H., & Epstein, R. A. (2016). The Occipital Place Area Is Causally Involved in Representing Environmental Boundaries during Navigation. Current Biology, 26(8), 1104–1109. 10.1016/j.cub.2016.02.066

33. Kaiser, D., Turini, J., & Cichy, R. M. (2019, October 9). A neural mechanism for contextualizing fragmented inputs during naturalistic vision. eLife; eLife Sciences Publications Limited. 10.7554/eLife.48182

34. Kamps, F. S., Julian, J. B., Kubilius, J., Kanwisher, N., & Dilks, D. D. (2016). The occipital place area represents the local elements of scenes. NeuroImage, 132, 417–424. 10.1016/j.neuroimage.2016.02.062

35. Kravitz, D. J., Peng, C. S., & Baker, C. I. (2011). Real-World Scene Representations in High-Level Visual Cortex: It’s the Spaces More Than the Places. Journal of Neuroscience, 31(20), 7322–7333. 10.1523/JNEUROSCI.4588-10.2011

36. Kupers, E. R., Knapen, T., Merriam, E. P., & Kay, K. N. (2024). Principles of intensive human neuroimaging. Trends in Neurosciences, 47(11), 856–864. 10.1016/j.tins.2024.09.011

37. Kwak, Y., & Curtis, C. E. (2022). Unveiling the abstract format of mnemonic representations. Neuron, 110(11), 1822–1828.e5. 10.1016/j.neuron.2022.03.016

38. Lescroart, M. D., & Gallant, J. L. (2019). Human Scene-Selective Areas Represent 3D Configurations of Surfaces. Neuron, 101(1), 178–192.e7. 10.1016/j.neuron.2018.11.004

39. Lever, C., Burton, S., Jeewajee, A., O’Keefe, J., & Burgess, N. (2009). Boundary vector cells in the subiculum of the hippocampal formation. Journal of Neuroscience, 29(31), 9771–9777.

40. Lin, T.-Y., Maire, M., Belongie, S., Hays, J., Perona, P., Ramanan, D., Dollár, P., & Zitnick, C. L. (2014). Microsoft coco: Common objects in context. Computer Vision–ECCV 2014: 13th European Conference, Zurich, Switzerland, September 6-12, 2014, Proceedings, Part v 13, 740–755.

41. Long, B., Yu, C.-P., & Konkle, T. (2018). Mid-level visual features underlie the high-level categorical organization of the ventral stream. Proceedings of the National Academy of Sciences, 115(38), E9015–E9024. 10.1073/pnas.1719616115

42. Mikolov, T., Chen, K., Corrado, G., & Dean, J. (2013). Efficient estimation of word representations in vector space. arXiv Preprint arXiv:1301.3781.

43. O’Keefe, J., & Dostrovsky, J. (1971). The hippocampus as a spatial map: Preliminary evidence from unit activity in the freely-moving rat. Brain Research, 34, 171–175. 10.1016/0006-8993(71)90358-1

44. Oliva, A., & Torralba, A. (2001). Modeling the shape of the scene: A holistic representation of the spatial envelope. International Journal of Computer Vision, 42, 145–175. 10/dm4d3b

45. Orghidan, R., Salvi, J., Gordan, M., & Orza, B. (2012). Camera calibration with two or three vanishing points.

46. Park, J., & Park, S. (2020). Coding of Navigational Distance and Functional Constraint of Boundaries in the Human Scene-Selective Cortex. The Journal of Neuroscience, 40(18), 3621–3630. 10.1523/JNEUROSCI.1991-19.2020

47. Park, S., Brady, T. F., Greene, M. R., & Oliva, A. (2011). Disentangling Scene Content from Spatial Boundary: Complementary Roles for the Parahippocampal Place Area and Lateral Occipital Complex in Representing Real-World Scenes. Journal of Neuroscience, 31(4), 1333–1340. 10.1523/JNEUROSCI.3885-10.2011

48. Park, S., Konkle, T., & Oliva, A. (2015). Parametric Coding of the Size and Clutter of Natural Scenes in the Human Brain. Cerebral Cortex, 25(7), 1792–1805. 10.1093/cercor/bht418

49. Pasternak, T., & Greenlee, M. W. (2005). Working memory in primate sensory systems. Nature Reviews Neuroscience, 6(2), 97–107.

50. Peer, M., & Epstein, R. A. (2021). The human brain uses spatial schemas to represent segmented environments. Current Biology, 31(21), 4677–4688.e8. 10.1016/j.cub.2021.08.012

51. Persichetti, A. S., & Dilks, D. D. (2018). Dissociable Neural Systems for Recognizing Places and Navigating through Them. The Journal of Neuroscience, 38(48), 10295–10304. 10.1523/JNEUROSCI.1200-18.2018

52. Portilla, J., & Simoncelli, E. P. (2000). A Parametric Texture Model Based on Joint Statistics of Complex Wavelet Coefficients. International Journal of Computer Vision, 40(1), 49–70. 10.1023/A:1026553619983

53. Prince, J. S., Charest, I., Kurzawski, J. W., Pyles, J. A., Tarr, M. J., & Kay, K. N. (2022). Improving the accuracy of single-trial fMRI response estimates using GLMsingle. eLife, 11, e77599. 10.7554/eLife.77599

54. Rademaker, R. L., Chunharas, C., & Serences, J. T. (2019). Coexisting representations of sensory and mnemonic information in human visual cortex. Nature Neuroscience, 22(8), Article 8. 10.1038/s41593-019-0428-x

55. Schubert, E., Rosenblatt, D., Eliby, D., Kashima, Y., Hogendoorn, H., & Bode, S. (2021). Decoding explicit and implicit representations of health and taste attributes of foods in the human brain. Neuropsychologia, 162, 108045. 10.1016/j.neuropsychologia.2021.108045

56. Serences, J. T., Ester, E. F., Vogel, E. K., & Awh, E. (2009). Stimulus-specific delay activity in human primary visual cortex. Psychological Science, 20(2), 207–214.

57. Smith, S. M., & Nichols, T. E. (2009). Threshold-free cluster enhancement: Addressing problems of smoothing, threshold dependence and localisation in cluster inference. NeuroImage, 44(1), 83–98. 10.1016/j.neuroimage.2008.03.061

58. Solstad, T., Boccara, C. N., Kropff, E., Moser, M.-B., & Moser, E. I. (2008). Representation of geometric borders in the entorhinal cortex. Science, 322(5909), 1865–1868.

59. Spelke, E. S., & Lee, S. A. (2012). Core systems of geometry in animal minds. Philosophical Transactions of the Royal Society B: Biological Sciences, 367(1603), 2784–2793. 10.1098/rstb.2012.0210

60. Wischnewski, M., & Peelen, M. V. (2021). Causal neural mechanisms of context-based object recognition. eLife, 10, e69736. 10.7554/eLife.69736

61. Wu, Y., & Li, S. (2024). Complexity matters: Normalization to prototypical viewpoint induces memory distortion along the vertical axis of scenes. Journal of Neuroscience, 44(27).

62. Xie, S., Kaiser, D., & Cichy, R. M. (2020). Visual Imagery and Perception Share Neural Representations in the Alpha Frequency Band. Current Biology, 30(13), 2621–2627.e5. 10.1016/j.cub.2020.04.074

63. Zhang, W., Zhang, W., & Zhang, Y. (2020). GeoLayout: Geometry Driven Room Layout Estimation Based on Depth Maps of Planes (No. arXiv:2008.06286). arXiv. http://arxiv.org/abs/2008.06286

